# An allele-sharing, moment-based estimator of global, population specific and population pairs *F*_*ST*_ under a general model of population structure

**DOI:** 10.1101/2023.07.18.549618

**Authors:** Jérôme Goudet, Bruce S. Weir

## Abstract

Being able to properly quantify genetic differentiation is key to understanding the evolutionary potential of a species. One central parameter in this context is *F*_*ST*_, the mean coancestry within populations relative to the mean coancestry between populations. Researchers have been estimating *F*_*ST*_ globally or between pairs of populations for a long time. More recently, it has been proposed to estimate population specific *F*_*ST*_ values, and between population-pair mean relative coancestry. Here, we review the several definitions and estimation methods of *F*_*ST*_, and stress that they provide values relative to a reference population. We show the good statistical properties of an allele sharing, method of moments based estimator of *F*_*ST*_ (global, population specific and population pairs) under a very general model of population structure. We point to the limitation of existing likelihood and Bayesian estimators when the populations are not independent. Last, we show that recent attempts to estimate absolute, rather than relative, mean coancestry fail to do so.

Author summary

## Introduction

Most species are spatially discontinuous. The partial isolation created by the discontinuity of the landscape limits gene flow. With time, genetic drift and possibly other random and non-random evolutionary processes will increase genetic differentiation among the partially isolated groups, which may favor or impede adaptation. Being able to properly quantify genetic differentiation is thus key to understanding the evolutionary potential of a species. Starting with the seminal work of Wright [1] and Malécot [2], population geneticists have been developing methods to this end.

*F*_*ST*_, a quantity very commonly estimated to measure genetic differentiation, is a key parameter, but there are multiple ways in which it can be defined. For instance, *F*_*ST*_ can be defined as the (intraclass) correlation between gametes chosen randomly from within the same population (the mean within populations coancestry) relative to the correlation for gametes chosen randomly from different populations (the mean between population coancestry) [3–6]. This amounts to using the studied set of populations as a reference point. Another definition uses the mean coancestry in an ancestral population as a reference point [4, 7, 8], which allows interpretation of *F*_*ST*_ in terms of identity by descent (IBD), but is valid only in very specific population models [4].

Several global estimators of *F*_*ST*_ have been developed, using the method of moments, maximum likelihood or Bayesian methods, as reviewed in [6]. These estimators have been used to quantify gene flow among populations [9], detect selection at specific loci [10, 11] or infer whether cooperation could evolve [12].

More recently, interest has turned to population and population-pair specific *F*_*ST*_ because of the realization that each population has a different history that can not be captured with a single average quantity [13–20].

The idea of population specific *F*_*ST*_ values was first proposed by Balding & Nichols [21] in a forensic context and was followed by a discussion of Bayesian estimation methods [22]. The authors used sample allele frequencies in a forensic database to produce posterior distributions of *F*_*ST*_ values for specific populations. They pointed to the greater benefit of using more loci, reducing the effects of genetic sampling variation, than of sampling more individuals and reducing the effects of statistical sampling.

Weir & Hill [13] provided method of moment estimators for population-specific and pairs of population *F*_*ST*_ values in work that made explicit mention of the dependence among populations of allele frequencies. They showed that their estimators were for within-population coancestry relative to average coancestry between pairs of populations. Population-specific estimates were shown to reveal differences at the LCT gene in humans, consistent with selection at that locus, that were not as evident with population-average estimates [23].

Beaumont & Balding [24] and Foll & Gaggiotti [25] also pointed to the use of population-specific *F*_*ST*_ ‘s in the context of identifying selected loci. Gaggiotti & Foll [26] reviewed the properties of a Bayesian estimator of population specific *F*_*ST*_ ‘s under the *F* model, which assumes all populations, possibly of different sizes, to descend from a single ancestral population and to receive immigrants from a common gene pool: a continent-islands model. These papers estimate allele frequencies in the ancestral population.

One limitation of estimators based on the *F* model is the assumption that the populations are independent [26]. Weir & Hill [13] relaxed that assumption, as did [14–16]. Karhunen & Ovaskainen [16] proposed a Bayesian estimator which allowed for specific migration rates between pairs of populations using an Admixture *F* model (AFM). Neither Weir & Hill [13] nor Karhunen & Ovaskainen [16] allowed for non random mating within populations, or provided predicted values in terms of demographic parameters. Mary-Huard & Balding [19] built on Ochoa & Storey [18] to develop a fast approximate method assuming a tree-like structure for the populations, but still assuming random mating (independence of gametes) within populations.

Weir & Goudet [17] provided estimators allowing for non random mating within populations, and gave predicted values of mean relative coancestries for a two-population system.

Most of this cited work did not investigate in depth the statistical properties of estimators. A notable exception is Gaggiotti & Foll [26] for population specific (but not population pair) *F*_*ST*_ that showed good qualitative properties of the Bayesian estimator of population specific *F*_*ST*_ under the *F* model, even with some departure from the model assumption of population independence. They did not provide predicted values for population specific *F*_*ST*_ other than for the island model.

Our goals here are fourfold. First we reiterate the well-known relation between *F*_*ST*_ and mean coancestries [3–5, 13, 17]. Second, we derive predicted values for coancestries and *F*_*ST*_ in a general model of population structure. Third, using computer simulations, we investigate the statistical properties of the moment based estimator of population specific and between population *F*_*ST*_ proposed by [17] using varying numbers of loci and individuals. We also evaluate in a subset of simulations the properties of the Bayesian estimator of *F*_*ST*_ put forth by [16]. Fourth, we present estimates for data from the 1000 Genomes project [27].

## Methods

### Definition of *F*_*ST*_

Rousset [4] gives a general definition of a parameter *F* as

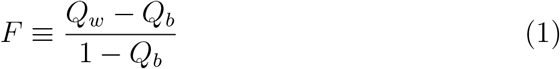

with *Q*_*w*_ and *Q*_*b*_ respectively defined as the probability of identity, or coancestry, within and between structural units.

He then states:

The well-known ‘F-statistics’ originally considered by Wright may be defined as above. […] For Wright’s *F*_*ST*_, *Q*_*w*_ is the probability of identity within a deme and *Q*_*b*_ is the probability of identity between demes. Likewise, Wright’s *F*_*IS*_, *Q*_*w*_ is the probability of identity of the two homologous genes in a diploid individual, and *Q*_*b*_ is the probability of identity of two genes in different individuals.

This general definition shows that *F*_*ST*_ and related quantities are always defined *relative to* a reference and that probabilities of identity themselves cannot be estimated. In what follows, we adopt the same framework.

### A general population model

We first present transition equations for mean coancestries between individuals, within and between populations in a metapopulation where populations can differ in effective size and exchange migrants according to a specified migration matrix. This general model includes as special cases (i) the continent-islands model, where islands receive alleles / immigrants from an infinite sized continent; (ii) the finite islands model, where all islands exchange migrants with all others at the same rate, as well as (iii) the stepping stone model, where populations exchange migrants only with their neighbors(see figure 1). Migration rates can either depend or not depend on the population sizes.

**Fig 1.**
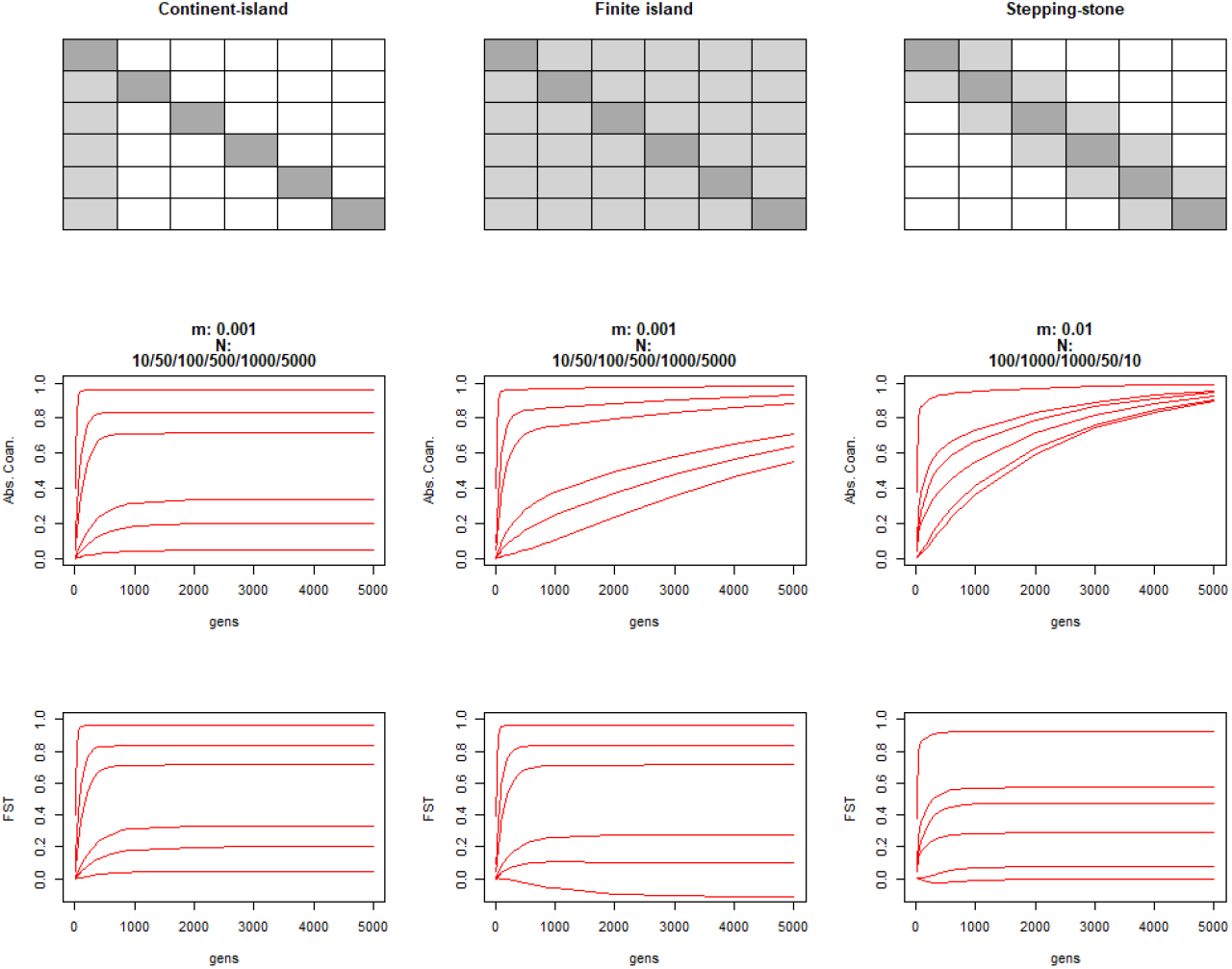
Migration, coancestries and *F*_*ST*_ for three migration models. The top row shows three migration matrices for sets of six populations, a continent-islands (left), where the continent (leftmost column) send migrants to all other columns, but receives none, and islands only receive immigrants from the continent; a Finite island (center) where each island send and received the same proportion to/from all other islands; and a finite one dimensional (1D) stepping stone model(right) where only the left and right neighbours contribute immigrants. White: cells with zero migration; light grey: cells with a positive migration term; dark grey: self. Middle row: Dynamics of within population coancestries through time, for the six different populations. Bottom row: Dynamics of population specific *F*_*ST*_ s through time, for the six different populations.

We assume a model where a number of populations of different effective sizes *N*_*i*_ are interconnected by migration, with a general migration matrix **M**. *N*_*i*_ are the effective rather than the census sizes, hence allowing approximately for any reproductive system. Elements of the matrix *m*_*ii*_*′*are the proportions of alleles in row population *i* that are from from column population *i*^*/*^ in the previous generation, including the case of *i* = *i*^*/*^. The only constraint on **M** is that all its elements are positive or zero, and each row sums to 1. The coancestries for distinct pairs of individuals in each population and between pairs of populations at time (*t* + 1) depend on their values at time *t* according to:

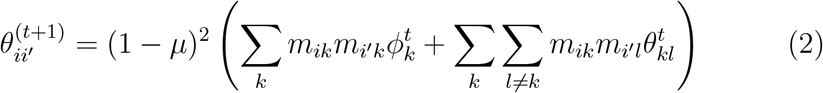

where 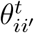 is the mean ibd probability for distinct pairs of individuals, one in population *i*, one in population *i*^*/*^, at time *t, µ* is the mutation rate, *ϕ*_*i*_ = 1*/*(2*N*_*i*_) + (2*N*_*i*_ − 1)*θ*_*i*_*/*(2*N*_*i*_) (we write *θ*_*i*_ or *θ*_*ii*_ for the mean ibd probability for distinct pairs of individuals both in population *i*) and *m*_*ii*_ = 1 − ∑ *i′*=*i m*_*ii*_*′*. Because *N*_*i*_ in the expression for *ϕ*_*i*_ is the effective size, where by definition mating occurs at random, any sex ratio bias or departure from random mating is accounted for, and since *θ*_*ii*_’s are for pairs of genes from different individuals, we need not worry about self coancestry. The model is valid for gametic migration and also for zygotic migration providing sampling occurs after dispersal. These equations can be rewritten in matrix form as:

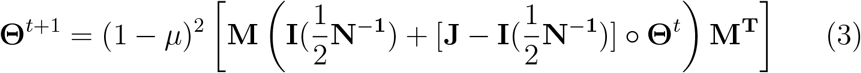

where **J** is a matrix of 1’s, **I** is the identity matrix, **M**^**T**^ is the transpose of **M, N**^−**1**^ is a vector with elements 1*/N*_*i*_ and ° denotes the Hadamard, or term by term, product of two matrices. We note this model can also accommodate extinction, fusion and fission of populations if we allow for the number of populations, effective populations sizes and migration rates to vary through time.

**Θ** in any generation contains the coancestries relative to those at time *t* = 0. To obtain ***F***_***ST***_, the matrix of coancestries *relative to* the mean coancestry for pairs of alleles, one in each of two different populations, we take the average, *θ*_*B*_, of the off-diagonal elements of **Θ**, subtract it from **Θ**, and divide the result by 1 − *θ*_*B*_:

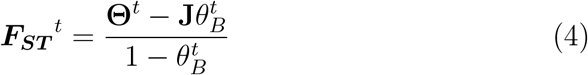

which has the same form as Equation 1. Averaging the diagonal elements of ***F***_***ST***_ over populations in any generation provides *F*_*ST*_ = (*θ*_*S*_ − *θ*_*B*_)*/*(1 − *θ*_*B*_) where *F*_*ST*_ is the overall *F*_*ST*_, and *θ*_*S*_, *θ*_*B*_ are the average coancestries within populations or between population pairs, respectively, for that generation. This was also given by [5, 13, 16, 17, 23]. The off-diagonal elements of ***F***_***ST***_ have an average zero by construction.

Eq 4 shows that *F*_*ST*_ for a pair of populations *i, i*^*/*^ involves all pairs of population-pair coancestries because of the *θ*_*B*_ term. The commonly used population pair *F*_*ST*_, however, consider only a particular pair of populations *i, i*^*/*^ and the quantity then being addressed is [(*θ*_*i*_ + *θ*_*i*_*′*)*/*2 − *θ*_*ii*_*′*]*/*(1 − *θ*_*ii*_*′*).

Classical population genetic models can be derived from this general one as follows:

- The continent-island model is obtained by setting the size of, e.g. the first population, to ∞, and allowing for migration terms only on the first column of the migration matrix (migration from the continent to the islands). The diagonal of **M** will thus consist of a 1 for the first element, and (1 − *m*_*i*1_) for the other diagonal elements, where *m*_*i*1_ is the proportion of immigrants in island *i* from the continent in the previous generation. For this model, eq. 3 can be rewritten:

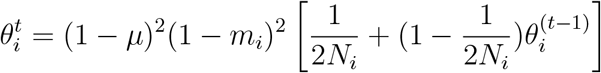

At equilibrium between mutation migration and drift, this gives (e.g. [3]):

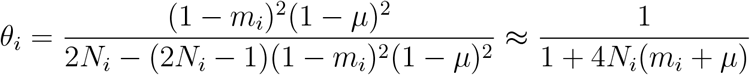

- The finite island model is obtained by setting all off-diagonal terms of the migration matrix to *m/*(*r* − 1), where *m* is the immigration rate and *r* the number of populations, and the diagonal elements to 1 − *m*.
- A 1 dimensional stepping stone model is obtained by setting the migration rate to 0 everywhere but to the left and right neighbor, whose rates are set to *m/*2, and on the diagonal, whose value is set to (1 − *m*). Populations at the left (or right) end of the set receive alleles only from the right (or left)

Analytical solutions for the dynamic of coancestries in the finite island model and the stepping stone model can be found in [28] and [29] respectively.

We illustrate with these three models how **Θ** and **F**_**ST**_ change through time for sets of six populations. Figure 1 shows the migration matrix and the dynamic of coancestries **Θ** through time for the continent island (left column), the finite island model (middle column) and the stepping stone model (right column). The populations all differ in effective sizes, varying from 10 to 5,000. The middle row shows the diagonal elements of **Θ**^*t*^ (the within population coancestries) while the bottom row shows the diagonal elements of **F**_**ST**_^*t*^ (the population specific *F*_*ST*_ ‘s) through time.

For the continent-islands model (left column), **Θ** (middle row) reach equilibrium before 1000 generations, and off diagonal elements are almost 0 (not shown), thus **F**_**ST**_ shows the same dynamic as mean coancestries. Small populations have larger mean coancestries and 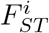 than large ones.

For the finite island model (middle column), we see a different pattern for **Θ**, as diagonal elements for the larger populations do not reach equilibrium within 5,000 generations, and the between population mean coancestries are also positive (not shown). On the other hand, all elements of **F**_**ST**_ have reached equilibrium, and we note that *F*_*ST*_ for the largest population is negative, implying a random pair of individuals from this population share less alleles on average than a random pair with one individual from one population and the second from another. The off-diagonal elements of **F**_**ST**_ also differ from 0, with elements between the largest populations being negative and those between small populations being positive (not shown).

For the 1-D stepping stone model (right column of Figure 1) we also see elements of the diagonal of **Θ** increasing through time, as do elements of the off diagonal elements (not shown), although at a slower pace. On the other hand, all elements of **F**_**ST**_ have reached an equilibrium value.

### A method of moments estimator for *F*_*ST*_

We now describe an allele sharing moment estimator of individual inbreeding, kinship and **F**_**ST**_ we first proposed in [17]. We allow for inbreeding (or any other form of non random mating, such as clonal reproduction) within populations. We index populations with superscripts *i, i*^*/*^, … and individuals with subscripts *j, j*^*/*^, The estimators are obtained from an *n*_*T*_ × *n*_*T*_ matrix **A** (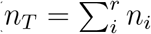, where *n*_i_ is the sample size for population *i*) of allele-sharing among individuals. For a bi-allelic diploid locus (see appendix S2 for a generalization to any ploidy level), allele-sharing between two individuals is 1 if the two individuals are homozygous for the same allele type, 0 if they are homozygous for different types, and 0.5 if at least one individual is heterozygous. Self-sharing is 1 for homozygous individuals and 0.5 for heterozygotes. When averaged over a large number of SNPs, this gives 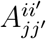, the allele sharing between individuals *j* and *j*^*/*^ in populations *i* and *i*^*/*^ respectively. Populations *i, i*^*/*^ and individuals *j, j*^*/*^ may or may not be the same. A moment estimator of self and between pairs of individuals kinship [30] is obtained as

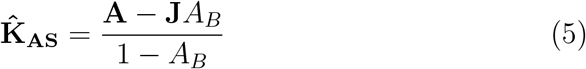

where *A*_*B*_ is the average of all the off-diagonal elements of **A** and **J** is a *n*_*T*_ × *n*_*T*_ matrix of 1s. Individual inbreeding coefficients relative to the total population are then estimated as 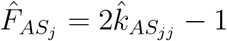 [31].

The mean allele sharing statistics between individuals within (*A*^*ii*^) and between (*A*^*ii*^*′*) population are obtained by averaging individual

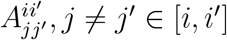 (thus self allele sharing is excluded from these averages) for each population *i* and population pair *i, i*^*/*^. These (*A*^*ii*^*′*) are stored in an *r* × *r* matrix **Ā**.

The individual inbreeding coefficient of individual *j* in population *i* 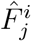, relative to the mean coancestry of population *i* is obtained as:

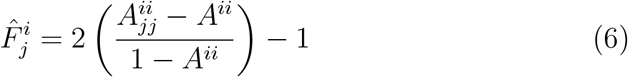

The average of the 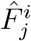s from one population gives an estimator of population *i* mean inbreeding coefficient 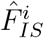. Averaging in turn these 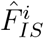 over populations lead to the overall estimator of within population inbreeding coefficient 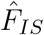, which, for equal sample sizes, is identical to Weir & Cockerham estimator of *F*_*IS*_ [32].

Writing as *ĀB* and *ĀS* the averages of the off-diagonal and diagonal elements of **Ā**, respectively, population specific *F*_*ST*_ estimates are obtained as:

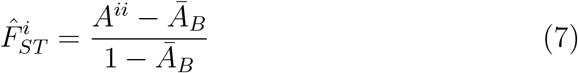

The average over populations of these 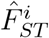 is an overall estimator of *F*_*ST*_ :

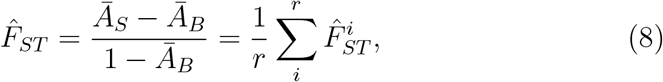

which, if all sample sizes are equal, is identical to Weir & Cockerham *F*_*ST*_ estimator for genotypes [32]

From matrix **Ā**, we can also obtain the following quantities:

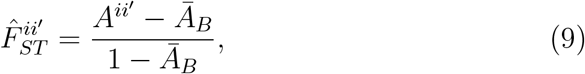

the mean coancestry for individuals, one in population *i* and one in population *i*^*/*^, relative to the average mean coancestry between pairs of populations. We are not aware of studies making use of the allele-sharing quantity described by eq. 9 although, for equal sample sizes, numerical values will be the same as those described by Weir & Hill [13]. Mary-Huard & Balding [19] described a similar quantity for tree like population structure, but did not explore its properties in detail.

For pairs of populations *i, i*^*/*^, using only those populations

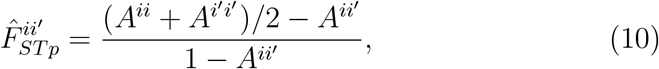

gives the average mean coancestry within populations *i* and *i*^*/*^ relative to the mean coancestry between populations *i* and *i*^*/*^. Eq. 10 gives the expression for pairwise *F*_*ST*_ often used to assess, for instance, whether isolation by distance is occurring in a dataset [20, 29].

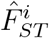 and 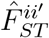 are conveniently stored in an *r* × *r* matrix 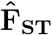:

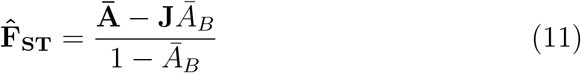

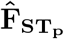, the matrix of pairwise *F*_*ST*_ values (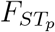, Eq. 10), can be retrieved instead from 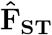 by replacing the *A*^*ii*^*′*s in Eq. 10 with the corresponding 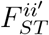 elements of 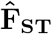.

While mean within and between population allele sharing **Ā** is not an unbiased estimator of mean within and between population coancestries **Θ** and *ĀB* is not an unbiased estimator of *θ*_*B*_, 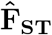 (Eq. 11) is an unbiased estimator of **F**_**ST**_ (Eq. 4) [17] as we will also show.

### Another moment estimator of F_ST_

Ochoa & Storey [18] recently proposed a closely related quantity. Rather than measuring average between-individual allele sharing within populations relative to average allele sharing between pairs of populations, they use the minimum average allele-sharing between populations as a reference, suggesting this should estimate ‘absolute mean coancestry relative to the most recent common ancestral population’ and claiming the measured quantities estimate ‘probabilities of Identity by Descent’. This new reference point changes the definition of ***F***_***ST***_ in Eq 4 to

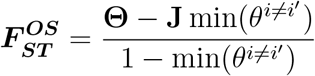

and we note that moving from one definition to the other is straightforward:

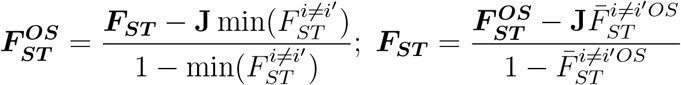

and can be extended to any constant one might want to use as a reference point (see appendix S1). The pairwise quantities estimated with Eq. 10 do not change with reference point.

### Statistical properties of 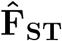

In order to investigate the statistical properties of population specific *F*_*ST*_ estimators, we first simulated genetic data sets with different population structures using the function `sim.genot.metapop.t` from the `R` package `hierfstat 0.5-11` [33, 34]. This function simulates independent loci from a series of populations connected by migration. Population can have different sizes and migration rates between populations is entered in matrix form, allowing among other things the simulation of the classical models of population structure described previously.

We focus on *F*_*ST*_ estimators for which the underlying model includes possible non independence of populations. This excludes estimators based on the strict *F* model [25, 26], as that model assumes all off-diagonal elements of **F**_**ST**_ to be zero. On the other hand, estimators based on the admixture *F* model (AFM, [16]) would be relevant.

The function `fs.dosage` from `hierfstat` was used to obtain the moment estimator 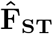. We used the `RAFM` package [35] and its `do.all` function with a burn-in of 5, 000 steps and a chain length of 10, 000, saving every fifth step to obtain the bayesian AFM coancestry estimates. The last 1, 000 saved steps from the MCMC were averaged to obtain the Bayesian estimates of **F**_**ST**_ using Eqs. 7 and 9. We checked in all cases the chain trace to insure it had converged. Predicted values for ***F***_***ST***_ are obtained using Eqs 3 and 4.

We use Root Mean Square Error (RMSE), the square root of the mean of squared deviations of the estimators from their expected values, to evaluate the different estimators and sampling strategies.

For simulations run with `sim.genot.metapop.t`, we estimated **F**_**ST**_ using the moment estimator with subsets of 1, 000 and 100 loci (100 replicates for each). For the Bayesian estimator, we only used 100 loci, as running time for more loci was prohibitive.

Nowadays most data sets consist of several thousand SNPs distributed randomly in the genome of the studied organisms and the genetic map length in a given species then becomes the ultimate constraint in the number of independent markers we can examine. In order to investigate the effects the genetic map length, the number of loci on the map and the number of individuals used, have on the bias and variance of 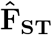, we ran a second set of simulations with the program `msprime` [36].

RMSE for data simulated with `msprime` were compared to the RMSE of data simulated with `sim.genot.metapop.t` with 10, 000 loci.

Finally, we estimated RMSE from the 1, 000 Genome data set too, using the values obtained from the complete data set as the expectation, and subsampling 10^5^, 10^4^, 10^3^, 5 × 10^2^ and 10^2^ SNPs per chromosome, and 50, 20, 10 or 5 individuals per population.

### Simulated population structure

We simulated three scenarios with different population structures. For each scenario, we simulated 20 replicates with 10, 000 loci, sampling 50 individuals using `sim.genot.metapop.t`.

- A finite island model with 10 islands with different sizes: *N*_1_ = *N*_2_ = 1000; *N*_3_ = *N*_4_ = 10; *N*_5_ = *N*_6_ = 100; *N*_7_ = *N*_8_ = 500; *N*_9_ = *N*_10_ = 2000, and a migration rate *m* = 0.001.
- A one dimensional stepping stone model with 10 populations, constant population sizes *N* = 1000 and *m* = 0.02 between adjacent populations.
- A river system with 2 tributaries rivers (see Figure 2 for details).

**Fig 2.**
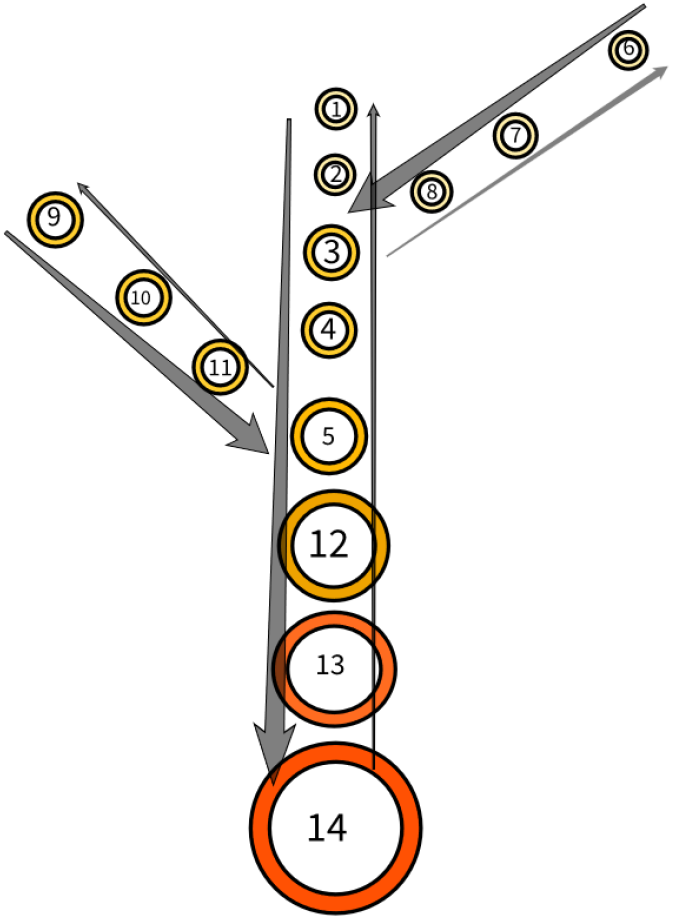
Sketch for the river system simulation. Each circle is a site, size and colour indicate population size. The main river is made of station 1 to 5 and 12 to 14. The first tributary river is made of stations 6 to 8 and joins the main river at station 3. The second tributary is made of stations 9 to 11 and joins the main river at station 5. Migration downstream to the nearest neighbor is *m*_*d*_ = 0.02 and migration upstream to the nearest neighbour is four times less, *m*_*u*_ = 0.005. Site 3 receives from and send migrants to sites 2 and 8, and site 5 receives from and sends migrants to sites 4 and 11. Population sizes increase as stations get closer to the mouth of the river, with *N*_1−2,6−8_ = 100; *N*_3−4,9−11_ = 200; *N*_5_ = 400; *N*_12_ = 800; *N*_13_ = 1, 000; *N*_14_ = 5, 000

We ran a second set of simulations for this river system structure with the program `msprime` [36]. For this second set of simulations, we generated five replicates of a genome made of 20 chromosomes each 10^8^ base pairs long with 4*Nc* = 4 × 10^3^ for each chromosome. This corresponds roughly to 10 centiMorgans (cM) in a population of 1, 000 individuals or 1 Morgan in a population of 10, 000 individuals.

## Results

### simulated data

**Figure 3** shows the results from the finite island model made of 10 populations with different sizes. The top left panel shows the expeected **F**_**ST**_. The smaller populations (3 and 4) show the largest 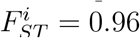 while the largest (9 and 10) have 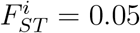. Off diagonal elements of **F**_**ST**_ are all fairly small between −0.06 and 0.09, those between large populations are slightly negative while those between small populations are positive. The top right panel shows RMSE distribution of 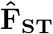 according to the sampling scheme or estimator used. We see a large effect of having fewer loci. The middle and bottom left panels illustrate that by using 10^3^ rather than 10^4^ SNPs, the variance of the estimates increases (but estimators remain unbiased), while subsampling individuals has almost no effect on the variance of the estimate (illustrated with the middle right panel where only 5 out of 50 individuals are sampled).

**Fig 3.**
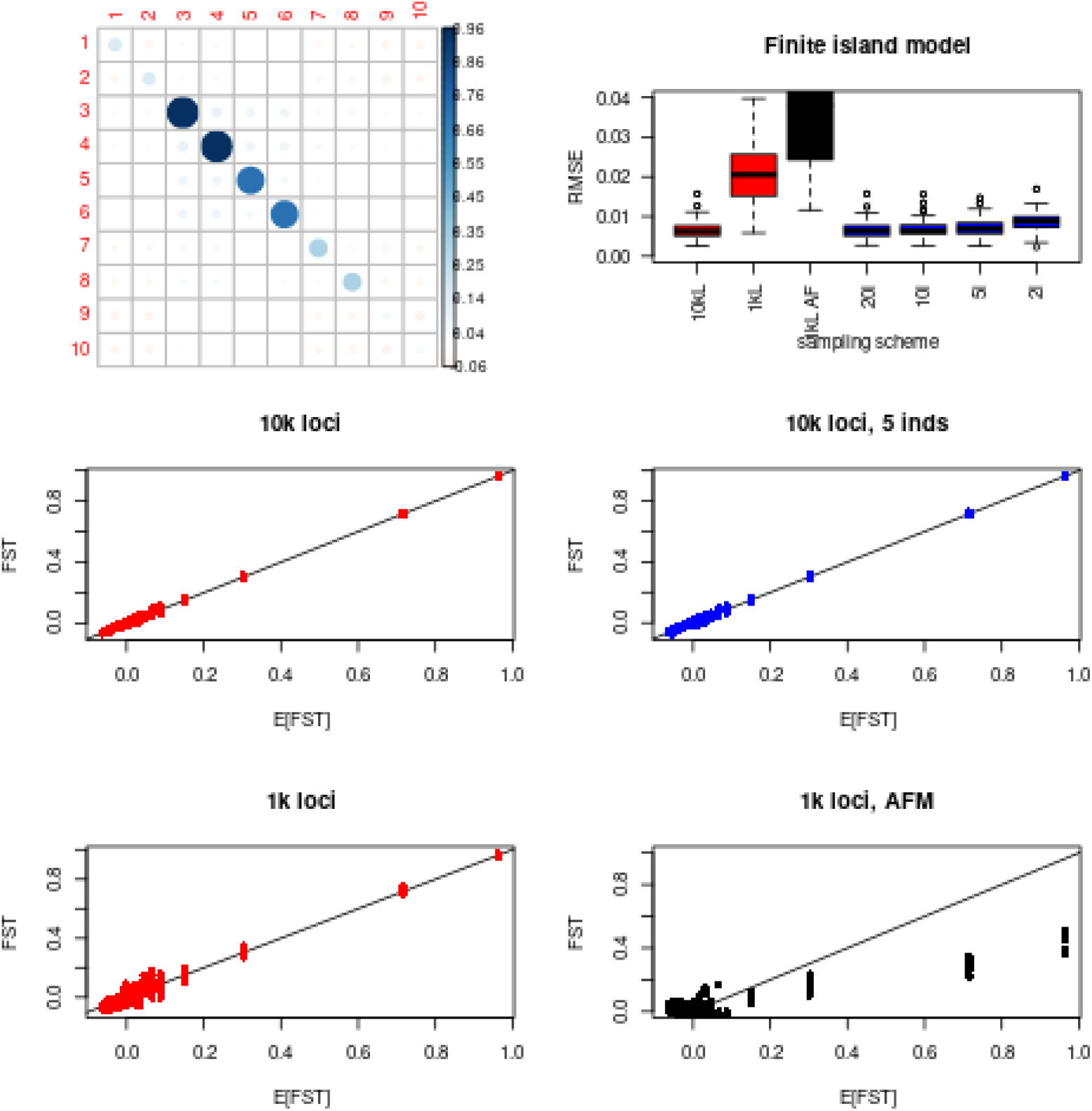
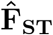 in a finite island model. 10 populations with different sizes: *N*_1_ = *N*_2_ = 1, 000; *N*_3_ = *N*_4_ = 10; *N*_5_ = *N*_6_ = 100; *N*_7_ = *N*_8_ = 500; *N*_9_ = *N*_10_ = 2, 000, and a migration rate *m* = 0.001. 50 individuals sampled in each population, with 10^4^ SNPs. Top left panel shows the expected (*F*_*ST*_) after 2, 000 generations from the transition equations (eq.4). Top right panel shows the distributions of Root Mean Square errors (RMSE) for all the elements of 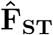 based on 20 or 100 replicates. ‘10kL, 1kL’: subsampling of 10^4^, 10^3^ SNPs respectively (red); ‘1kL AFM’: 1, 000 SNPs, using the Bayesian estimator from AFM (black); ‘20i, 10i, 5i, 2i’: Subsampling of 20, 10, 5 or 2 individuals, 10^4^ SNPs (blue). The four lower panels show the relation between expected and estimated *F*_*ST*_ s for 10^4^ SNPs (middle left), 10^4^ SNPs, 5 individuals (middle right), 1k SNPs (bottom left), 1k SNPs, AFM method (bottom right), with red showing subsampling of loci, blue subsampling of individuals and black the bayesian estimate

The Bayesian estimates (bottom right panel) show large biases, with an underestimation of large values and an overestimation of low values of the elements of **F**_**ST**_.

**Figure 4** shows results for the one dimensional stepping stone model. The top left panel shows **F**_**ST**_. Populations at the end of the stepping shows the largest 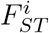 and those at the centre positive but low values. Off-diagonal elements of populations far apart are negative, the further apart the more negative.

**Fig 4.**
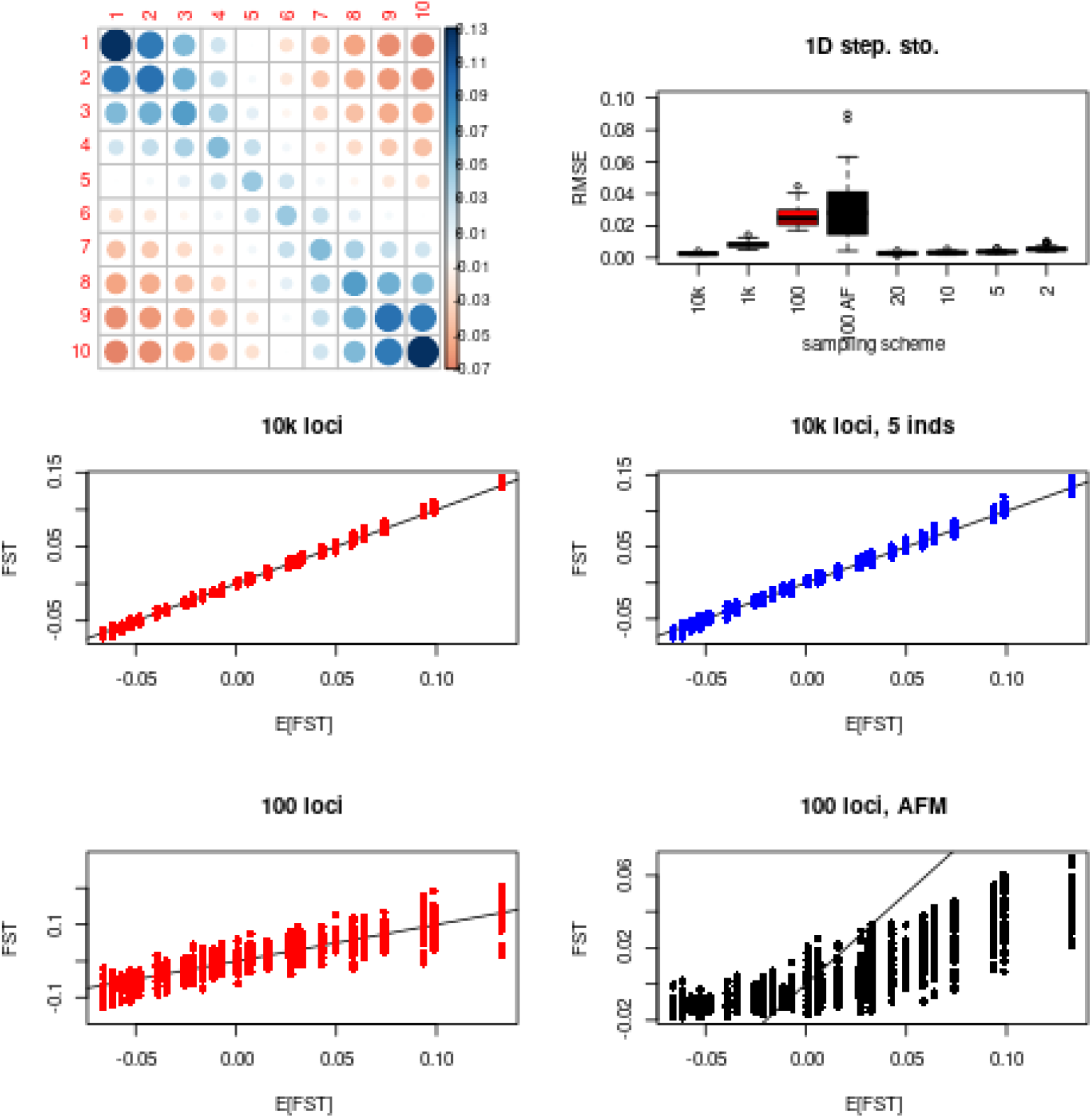
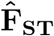 in a 1D stepping stone model. 10 populations with *N* = 1, 000; *m* = 0.005 between adjacent populations. 50 individuals sampled in each population, with 10^4^ SNPs. Top left panel shows the expected (*F*_*ST*_) after 4, 000 generations from the transition equations (eq.4). Top right panel shows the distributions of Root Mean Square errors (RMSE) for all the elements of 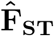 based on 20 or 100 replicates. ‘10k, 1k, 100’: 10^4^, 10^3^ and 100 SNPs respectively (red); ‘100 AFM’: 100 loci, using the bayesian estimator from `AFM` (black); ‘20, 10, 5, 2’: Subsampling of 20, 10, 5 or 2 individuals, 10^4^ loci (blue). The four lower panels show the relation between expected and estimated *F*_*ST*_ s for 10^4^ SNPs (middle left), 10^4^ SNPs, five individuals (middle right), 100 SNPs (bottom left), 100 SNPs, AFM method (bottom right), with red showing subsampling of loci, blue subsampling of individuals and black the bayesian estimate than other elements, and negative elements appear for pairs where one member is close to the mouth and the other is in one of the tributaries.

The top right panel shows RMSEs of 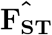 for varying number of loci and individuals. We see the same pattern as was seen for the island model: the fewer loci, the larger are RMSE, but with no bias for the moment estimator (middle and bottom left panels). Sub-sampling five individuals per population out of 50 has almost no effect on RMSE values (middle right panel). The Bayesian estimates are strongly biased, with overestimation for low values of the parameter and underestimation for high values.

**Figure 5** shows results for the river system, where populations differ in size and migration is asymmetric. The top left panel shows **F**_**ST**_. The largest 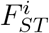 are for stations 1 and 6, those close to the sources of the river system, while the smallest is for population 14, at the mouth of the river. The off-diagonal elements of the tributaries further up the river are larger

**Fig 5.**
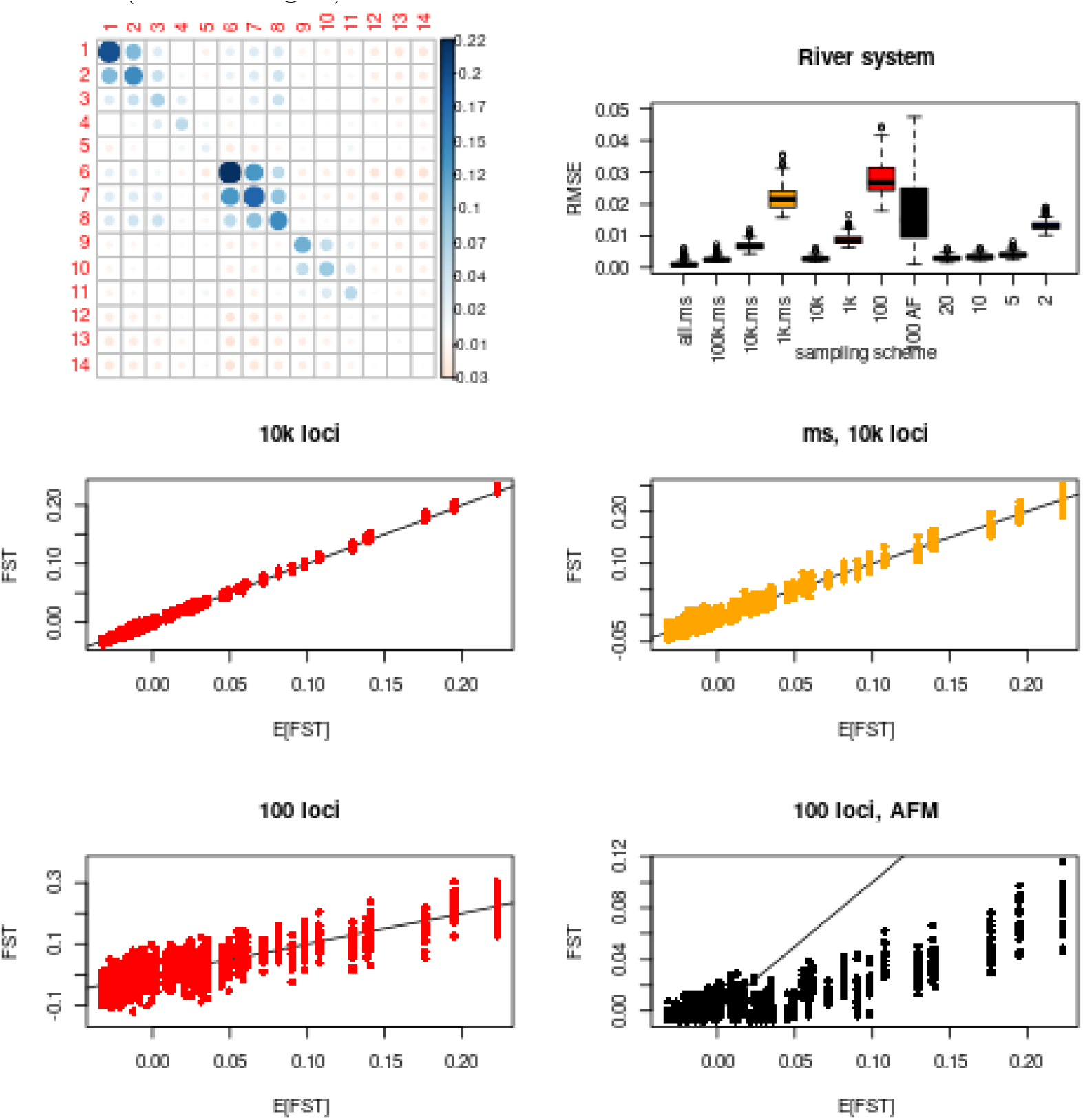
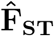 in a river system. see methods for description of the system. 50 individuals sampled in each population, with 10^4^ SNPs. Top left panel shows the expected (*F*_*ST*_) after 2, 000 generations from the transition equations (eq.4). Top right panel shows the distributions of Root Mean Square errors (RMSE) for all the elements of 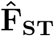 based on 20 or 100 replicates. ‘all.ms, 100k.ms, 10k.ms, 1k.ms’: All, 10^5^, 10^4^, 10^3^ SNPs from `msprime` simulations (yellow); ‘10k, 1k, 100’: 10^4^, 10^3^ and 100 SNPs respectively (red); ‘100 AFM’: 100 loci, using the bayesian estimator from `AFM` (black); ‘20, 10, 5, 2’: Subsampling of 20, 10, 5 or 2 individuals, 10^4^ loci (blue). The four lower panels show the relation between expected and estimated *F*_*ST*_ s for 10^4^ SNPs (middle left), 10^4^ SNPs from `msprime` simulations (middle right), 100 SNPs (bottom left), 100 SNPs, AFM method (bottom right)

The effect of subsampling independent loci is similar to what we have seen for the island and stepping stone model, with RMSEs increasing with fewer loci (red boxplots, top right panel, and middle and bottom left panels for 10^4^ and 100 SNPs respectively). Subsampling individuals has little effect on RMSEs but for 2 individuals per population, where RMSEs increases.

The Bayesian estimator again give biased estimates, with an underestimation of large and an overestimation of small elements (bottom right panel).

Finally for this population structure, we also simulated data with `msprime` to look at the effect of linkage among SNPs (orange boxplot in top right panel). Compared with the simulations with independent SNPs, roughly 10 times more SNPs are necessary to obtain similar RMSEs. The graphs on the middle row of the figure shows the precision of the estimates for 10^4^ SNPs, with independent SNPs on the left and partially linked SNPs on the right, where the variation is larger than with independent SNPs.

### Estimates of F_ST_ in the 1000 genomes

**Figure 6** shows estimates of **F**_**ST**_ and pairwise 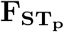 in the 1000 genomes. Looking at the top left panel (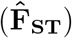) we see all estimates where at least one of the populations is from Africa are negative, while all other estimates are positive. A negative value of 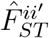 implies that the pairs of populations considered have less allele sharing than random pairs from the whole world. As African genomes are highly heterozygous, it is not surprising they show the lowest allele sharing with other African (including those from their population) or non African genomes. Populations from East Asia shows the largest 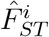 values, followed by European and South Asians. Admixed American populations are the more heterogeneous, with Puerto Ricans from Puerto Rico (PUR) showing the lowest values and Peruvian from Lima (PEL) the highest (Supplementary Figure S1).

**Fig 6.**
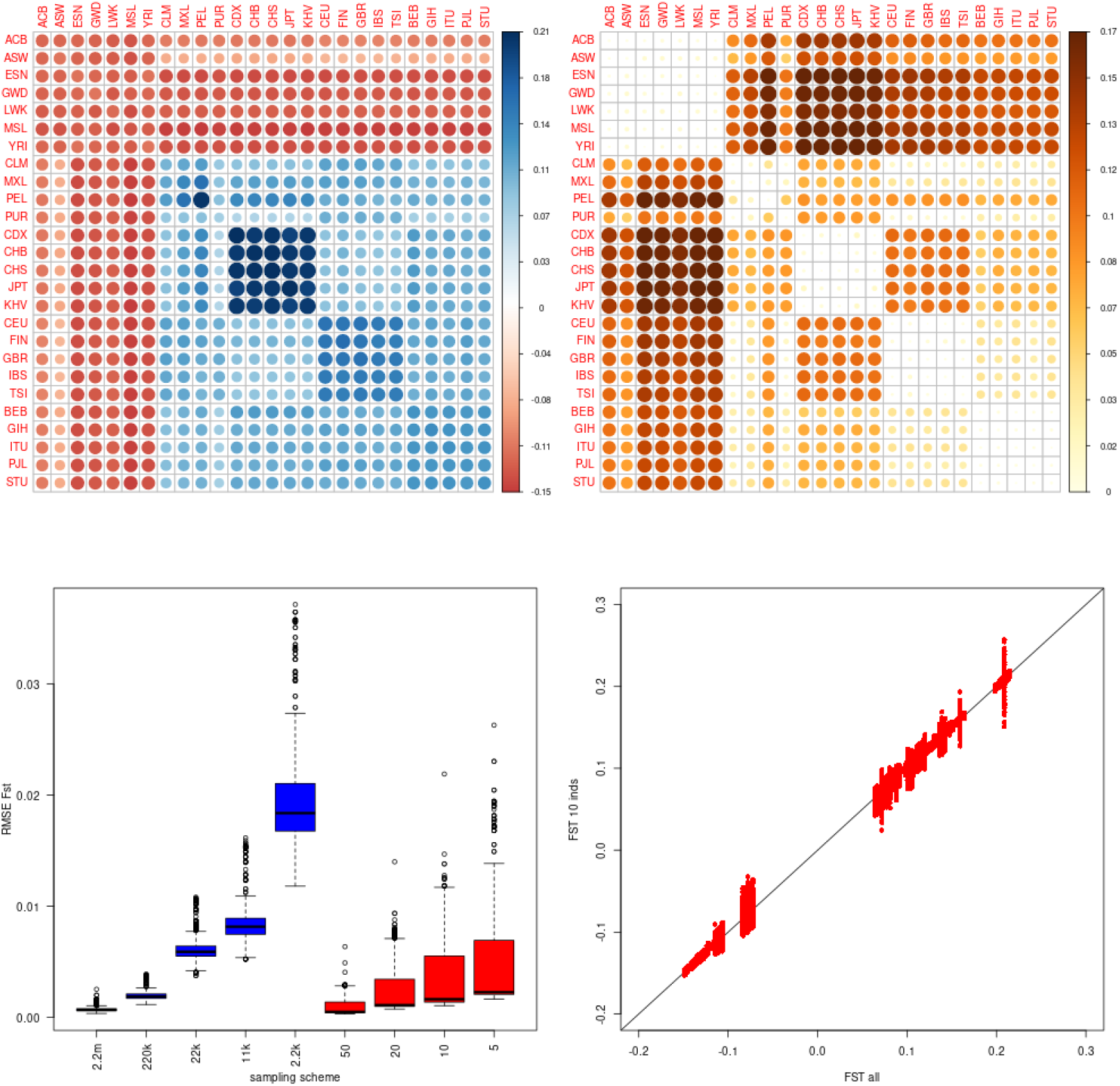
estimates of F_ST_ from the 1000 genomes. The top row shows 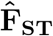 (left), and pairwise 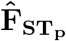 (right) estimated from the whole 1000 genome dataset, with color scale to the right of each panel. The bottom, left panel shows the distribution of RMSEs for 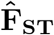. ‘2.2m, 220k, 22k, 2.2k’ (in blue): subsampling of the corresponding number of SNPs from the total data set; ‘50, 20, 10, 5’ (in red); subsampling of the corresponding number of individuals from the total dataset. The bottom right panel shows 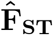 estimated from 10 individuals (100 replicates) against 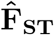 estimated from all individuals.

The top right panel of Figure 6 shows pairwise 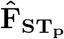. Here all values are positive (a property of 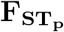), and all values for pairs of populations from the same continent are close to 0. African populations show the largest estimates with populations from other continents, in particular East Asia and PEL. Among non African populations, the largest differences are between East Asian and European samples.

The bottom left panel shows RMSE for subsampling loci in blue and individuals in red. Using fewer loci and individuals increases RMSE for 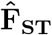.

The effect of subsampling individuals on RMSE for 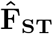 differs from what we have seen in the simulations, as we observed some elements of the RMSE for 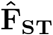 have larger values (Figure S2). From the bottom right panel of Figure 6, we see as few as 10 individuals could give results almost identical to 100 in homogeneous populations, but this is not so for admixed populations, where we see large variation among replicates when we subsample 10 individuals.

## Discussion

In this paper, we provide predicted values for **F**_**ST**_, the mean coancestries within and among populations relative to the average between populations mean coancestries, for a very general model of population structure in a diploid species, and show our allele sharing, moment based estimator of **F**_**ST**_ to be efficient and unbiased in all simulation scenarios, and for all elements (on and off-diagonal) of the matrix. The Bayesian estimator [16] is biased for all simulated scenarios and does not scale to the size of genomic datasets generated nowadays.

We find that as few as 10 or even five individuals per population are sufficient to estimate accurately **F**_**ST**_, unless the population contains admixed individuals. With 10^4^ SNPs, estimates are very accurate, and 10^3^ independent SNPs might actually be enough. However, one should be aware that if SNPs are tightly linked, as may be the case with small genomes and/or small populations size (the relevant quantity is 4*Nr*, where *N* is the effective size and *r* the recombination fraction between SNPs), more SNPs are needed, as we illustrate with the river system example.

In real data, it is unclear how linkage should be accounted for. In practice, either blocks of a constant number of SNPs or blocks of a constant number of nucleotides are used for bootstrapping, but it might be more appropriate to define block size according to the recombination rate of the different regions, and we note this is an area of active research [37]. Here, we obtain confidence intervals for continent and overall 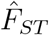 in the 1000 genomes by bootstrapping estimates obtained from blocks of 100 kilobases, but recognized that this is *adhoc* and an avenue for further research.

In deriving 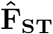, we make no assumptions about mating system, inbreeding level, ploidy level or even whether reproduction is sexual [17]. As long as allele sharing between individuals can be measured, 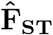 can be estimated.

### Use of 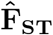

Global *F*_*ST*_ and pairwise 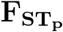 are commonly estimated in surveys of population structure, and population specific *F*_*ST*_ ‘s have also been used [23, 26, 38]. But we are not aware of studies estimating the off-diagonal elements of **F**_**ST**_.

One area where such measures are useful is in conservation genetics, as the off-diagonal elements inform about how much of the overall genetic diversity is captured by the pair of populations considered. Large positive values indicate that the two populations represent a small proportion of the overall diversity, while low and negative values indicate to the contrary that these two populations together harbour as much as or more diversity than all populations together. For instance, in the stepping stone example (Figure 4), populations 1 and 10 each contain less variability than others (large 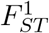 and 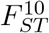), as can be seen from the diagonal elements of the top left panel, but are also those that together harbour the most diversity (lowest off-diagonal element is *F* ^1,10^). In the river system example (Figure 5), the populations harbouring the most diversity together (those with the lowest 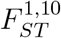) would be 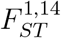 and 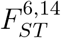, but we see that the pair 1 and 6, with the largest pairwise 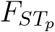 (not shown), does not have the lowest *F*_*ST*_, and 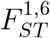 is positive.

The size of 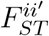 does not inform about the size of 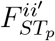: imagine two populations have fixed the same allele over the majority of the SNPs, but fixed opposite one at a few loci. 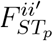 would be one, but depending on the genetic make up of other populations, 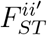 could be very large, meaning these two populations together capture only a small fraction of the overall diversity. On the other hand, if the two populations have fixed alternate alleles at most of their loci, 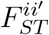 would still be one but 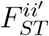 is likely going to be negative, as these two populations together would show maximum possible diversity.

### Choice of a reference point

Ochoa and Storey [18] suggest using the allele sharing between the least related pair of populations, rather than the average between population allele sharing, as a reference point. They claim this population pair with the lowest allele sharing represents the closest to what would have been the ancestral population. By using the minimum between population allele sharing as a reference point, they insure that all terms of 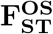 are positive, because they want to interpret these values in term of identity by descent. But we note their estimates are not probabilities of identity by descent.

Karhunen & Ovaskainen [16] also estimate coancestries relative to an ancestral population, but the model they use, the admixture F model (AFM), is more restrictive than ours or Ochoa & Storey’s [18], and they were careful in distinguishing between mean coancestries, estimated relative to the allele frequencies in the ancestral gene pool (see figure 1 and Eq. 12 in [16] and *F*_*ST*_ (Eq. 4 in [16]). While we see great values in obtaining estimates of mean coancestries relative to a reference population in the past (for instance in the context of detecting local adaptation on traits, see [39, 40]), we emphasized these coancestries differ from the standard definition of *F*_*ST*_ given in Equation 1 [5, 16, 17, 41].

While the algebraic difference in the formulae between our estimator and that proposed by Ochoa & Storey [18] is trivial, we show with data from the 1000 Genomes project (Table S3 and Figure S3) that the two estimates can greatly differ (Overall 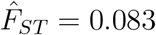 against overall 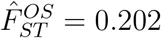). While 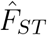 is similar to previously reported *F*_*ST*_ for human populations, values as large as 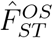 have not been reported. It is interesting to note that the text book estimator of *F*_*ST*_, 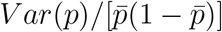, which, with these samples sizes (2.504 individuals, ≈ 100 individuals per population and 26 populations), should be little affected by statistical biases, gives a value for the same dataset of 0.088 (range [0.083, 0.094]), much closer to 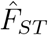 than to 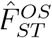.

Two other considerations indicate that 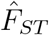 is to be preferred:

When comparing 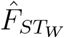 and 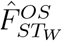 in the 1000 genomes, we see the ranks among chromosomes is not conserved (Table S3 and Figure S3), because while the transformation between the two estimators is linear, each chromosome will show a different minimum value (and a different arg min value), and hence the linear transformation for each chromosome is different. For example, chromosome 21 is second lowest with 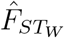 but twelfth lowest with 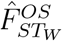. We also find that the confidence intervals for are 2.25 to 3.5 times wider than those for 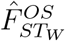 (Figure S3).

Imagine a very large number of populations. All but two populations (1 and 2) contains only heterozygotes at all loci (we would obtain the same result with any genotypic composition maintaining an allelic frequency at 0.5 in all populations but the first 2), and populations 1 and 2 are fixed for the opposite homozygotes at all loci. For populations 1 and 2, fixed for opposite homozygotes, 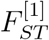 and 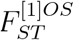 will be one, as will 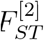 and 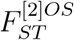. 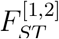 will tend to −1, while 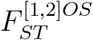 will be 0, by definition. All other 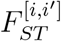 will tend to ∞ as the number of populations tend to, but will tend to 0.5 for 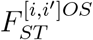. We believe a value of 0 is more meaningful than 0.5 for these pairs with identical genotypic composition.

## Conclusions

We showed that a moment estimator of **F**_**ST**_ based on allele sharing and not making any assumptions about population independence, mating system, ploidy level or inbreeding status of individuals, is unbiased, accurate, fast to calculate and scales easily to genome size data. We provide numerical and analytical solutions for the expectation of **F**_**ST**_ given migration and mutation rates and population sizes allowing investigators to evaluate how these parameters will affect **F**_**ST**_. The function `fs.dosage` from the `hierfstat (0.5-11)` R package [33, 34] estimates **F**_**ST**_ and 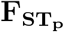, as well as individual inbreeding coefficients.

## Supporting information

### S1 A general reference point for kinship and *F*_*ST*_

In general

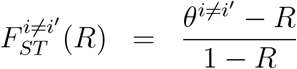

where we take *R* = mean(*θ*^*i* ≠*i*^′) and OS take *R* = min(*θ*^*i*≠ *i*^′). For two different values *R, R*^*/*^:

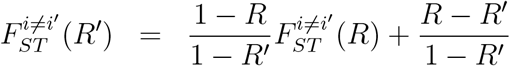

### S2 Allele sharing, kinship and inbreeding for a *k*−ploid species

for a *k* ploid species, given a dosage matrix **X** with *n* rows (individuals) and *L* columns (loci) and with element [*j, l*] corresponding to the number of copies of the designated allele for the *j*th individual at the *l*th locus, hence taking (integer) values between 0 and *k*, allele sharing between the different possible dosages at a locus for that ploidy level are given in table S1. The allele sharing matrix 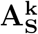 is obtained by counting for each pair of individuals the proportion of loci in one of the *k*^2^ states given in table S1 and multiplying this proportion by the corresponding allele sharing. This is efficiently done using the following:

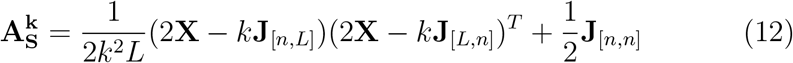

where **J**_[*a,b*]_ is a matrix of ones with *a* rows and *b* columns. An equivalent expression was independently derived by Bilton [42]

calling *A*_*B*_ the average of all the off-diagonal elements of 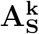, kinships are obtained as

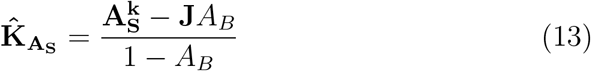

With the inbreeding coefficient for a *k* − ploid individual defined as the probablility that 2 alleles drawn at random without replacement from the *k* the individual carries are identical (self kinship being the probability that 2 alleles drawn at random with replacement are identical), individual inbreeding coefficients are obtained from self kinship as:

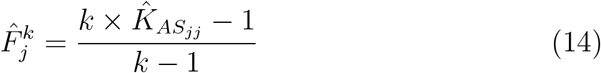

which reduces to the classical 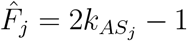 for a diploid species (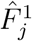 is of course undefined for haploid species).

**Table S1.**
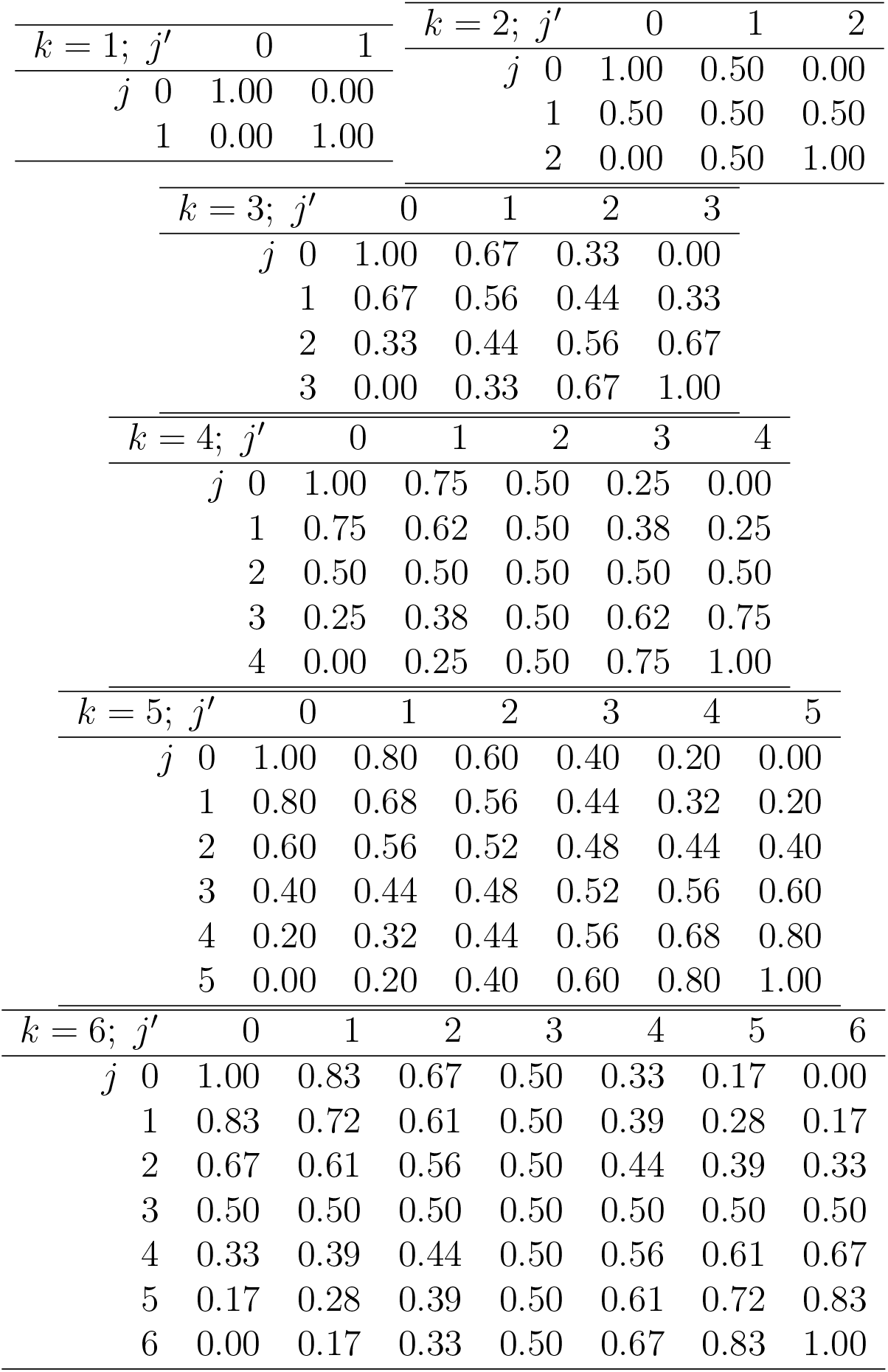
Allele sharing probabilities *A*_*jj*_*′*between the different possible dosages for ploidies *k* from 1 to 6

### S3 One thousand genomes population statistics

Table S2 shows continent-specific and figure S1 shows population-specific estimates of *F*_*ST*_ obtained from phase 3 data of the 1000 genomes project.

---

**Fig S1.**
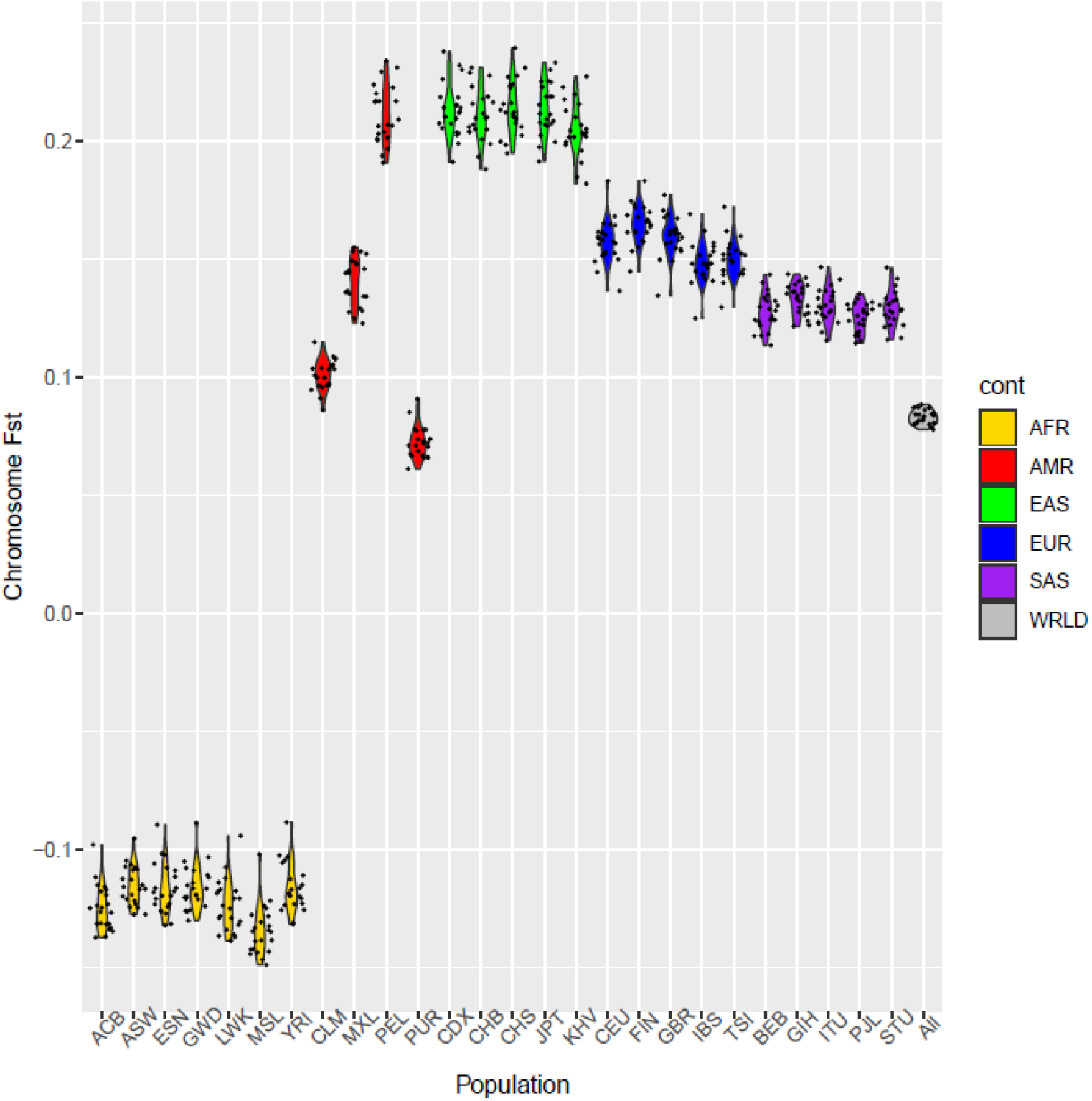
Population specific estimates of *F*_*ST*_ from the 1000 genomes. Violin plots of the chromosome level, population specific *F*_*ST*_, using data from phase 3 of the 1000 genomes project. Populations are ordered by continent, Africa in gold, America in red, East Asia in green, Europe in blue and South Asia in purple. Average overall all populations is shown in grey. Each dot represents the estimate obtained from one of the 22 autosomes.

**Fig S2.**
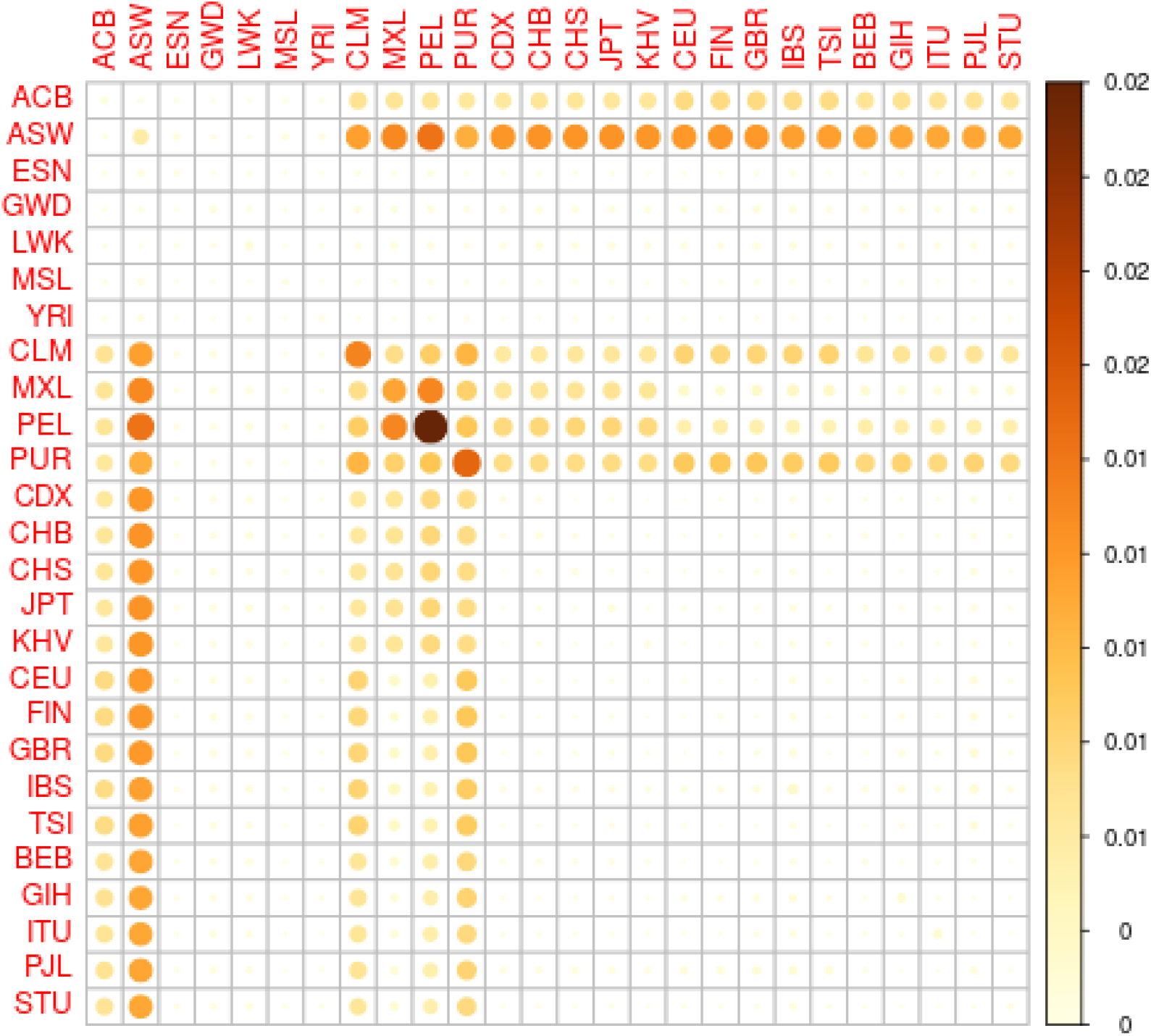
RMSEs of 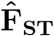 in the 1000 genomes when subsampling 10 individuals per population. Color scale on the right hand side of the figure. The larger the size of the circle and the darker the color, the higher the RMSEs for this population/ population pair. All the populations that stand out are the admixed ones.

**Table S2.** Estimates of chromosome and continent specific *F*_*ST*_ from the 1000 genomes and their 95% confidence intervals obtained by block bootstrap, using 100kb blocks

| Chrom. | AFR | AMR | EAS | EUR | SAS |
| --- | --- | --- | --- | --- | --- |
| 1 | -0.137<br>(-0.145, -0.131) | 0.100<br>(0.097, 0.103) | 0.198<br>(0.193, 0.204) | 0.152<br>(0.147, 0.157) | 0.120<br>(0.116, 0.123) |
| 2 | -0.145<br>(-0.151, -0.137) | 0.102<br>(0.099, 0.106) | 0.217<br>(0.211, 0.222) | 0.146<br>(0.142, 0.151) | 0.126<br>(0.122, 0.130) |
| 3 | -0.140<br>(-0.147, -0.133) | 0.096<br>(0.093, 0.099) | 0.205<br>(0.198, 0.211) | 0.144<br>(0.140, 0.149) | 0.121<br>(0.118, 0.126) |
| 4 | -0.138<br>(-0.145, -0.132) | 0.101<br>(0.097, 0.105) | 0.197<br>(0.192, 0.205) | 0.146<br>(0.141, 0.150) | 0.117<br>(0.113, 0.121) |
| 5 | -0.151<br>(-0.157, -0.144) | 0.097<br>(0.094, 0.100) | 0.207<br>(0.202, 0.214) | 0.141<br>(0.135, 0.145) | 0.121<br>(0.116, 0.125) |
| 6 | -0.109<br>(-0.118, -0.102) | 0.091<br>(0.086, 0.094) | 0.180<br>(0.172, 0.188) | 0.123<br>(0.119, 0.130) | 0.107<br>(0.103, 0.112) |
| 7 | -0.133<br>(-0.142, -0.128) | 0.096<br>(0.092, 0.099) | 0.192<br>(0.186, 0.200) | 0.140<br>(0.135, 0.144) | 0.115<br>(0.111, 0.120) |
| 8 | -0.152<br>(-0.161, -0.144) | 0.103<br>(0.100, 0.107) | 0.206<br>(0.199, 0.213) | 0.155<br>(0.149, 0.161) | 0.119<br>(0.116, 0.124) |
| 9 | -0.137<br>(-0.146, -0.127) | 0.102<br>(0.099, 0.107) | 0.193<br>(0.184, 0.200) | 0.147<br>(0.141, 0.153) | 0.113<br>(0.107, 0.118) |
| 10 | -0.123<br>(-0.132, -0.116) | 0.093<br>(0.089, 0.097) | 0.196<br>(0.188, 0.205) | 0.140<br>(0.135, 0.148) | 0.111<br>(0.107, 0.115) |
| 11 | -0.137<br>(-0.145, -0.132) | 0.100<br>(0.096, 0.103) | 0.195<br>(0.188, 0.204) | 0.134<br>(0.129, 0.140) | 0.121<br>(0.116, 0.124) |
| 12 | -0.126<br>(-0.134, -0.118) | 0.100<br>(0.096, 0.103) | 0.196<br>(0.188, 0.206) | 0.141<br>(0.135, 0.146) | 0.118<br>(0.113, 0.124) |
| 13 | -0.130<br>(-0.138, -0.120) | 0.096<br>(0.091, 0.101) | 0.196<br>(0.185, 0.204) | 0.136<br>(0.130, 0.142) | 0.114<br>(0.108, 0.118) |
| 14 | -0.129<br>(-0.137, -0.120) | 0.108<br>(0.103, 0.113) | 0.184<br>(0.176, 0.192) | 0.152<br>(0.144, 0.159) | 0.116<br>(0.111, 0.122) |
| 15 | -0.145<br>(-0.157, -0.136) | 0.108<br>(0.104, 0.114) | 0.196<br>(0.190, 0.206) | 0.165<br>(0.157, 0.173) | 0.123<br>(0.116, 0.130) |
| 16 | -0.153<br>(-0.163, -0.140) | 0.107<br>(0.103, 0.112) | 0.200<br>(0.191, 0.209) | 0.151<br>(0.143, 0.157) | 0.124<br>(0.118, 0.130) |
| 17 | -0.145<br>(-0.155, -0.134) | 0.102<br>(0.094, 0.108) | 0.221<br>(0.211, 0.232) | 0.141<br>(0.131, 0.151) | 0.132<br>(0.125, 0.138) |
| 18 | -0.142<br>(-0.155, -0.131) | 0.094<br>(0.089, 0.099) | 0.188<br>(0.179, 0.196) | 0.143<br>(0.135, 0.150) | 0.123<br>(0.117, 0.128) |
| 19 | -0.148<br>(-0.160, -0.137) | 0.101<br>(0.096, 0.106) | 0.202<br>(0.191, 0.211) | 0.145<br>(0.138, 0.154) | 0.108<br>(0.103, 0.115) |
| 20 | -0.149<br>(-0.161, -0.135) | 0.101<br>(0.096, 0.107) | 0.212<br>(0.203, 0.225) | 0.145<br>(0.139, 0.152) | 0.128<br>(0.122, 0.135) |
| 21 | -0.146<br>(-0.161, -0.131) | 0.108<br>(0.101, 0.115) | 0.181<br>(0.172, 0.193) | 0.143<br>(0.134, 0.153) | 0.114<br>(0.104, 0.123) |
| 22 | -0.128<br>(-0.142, -0.112) | 0.109<br>(0.103, 0.115) | 0.214<br>(0.197, 0.227) | 0.138<br>(0.130, 0.145) | 0.111<br>(0.102, 0.119) |

**Table S3.**
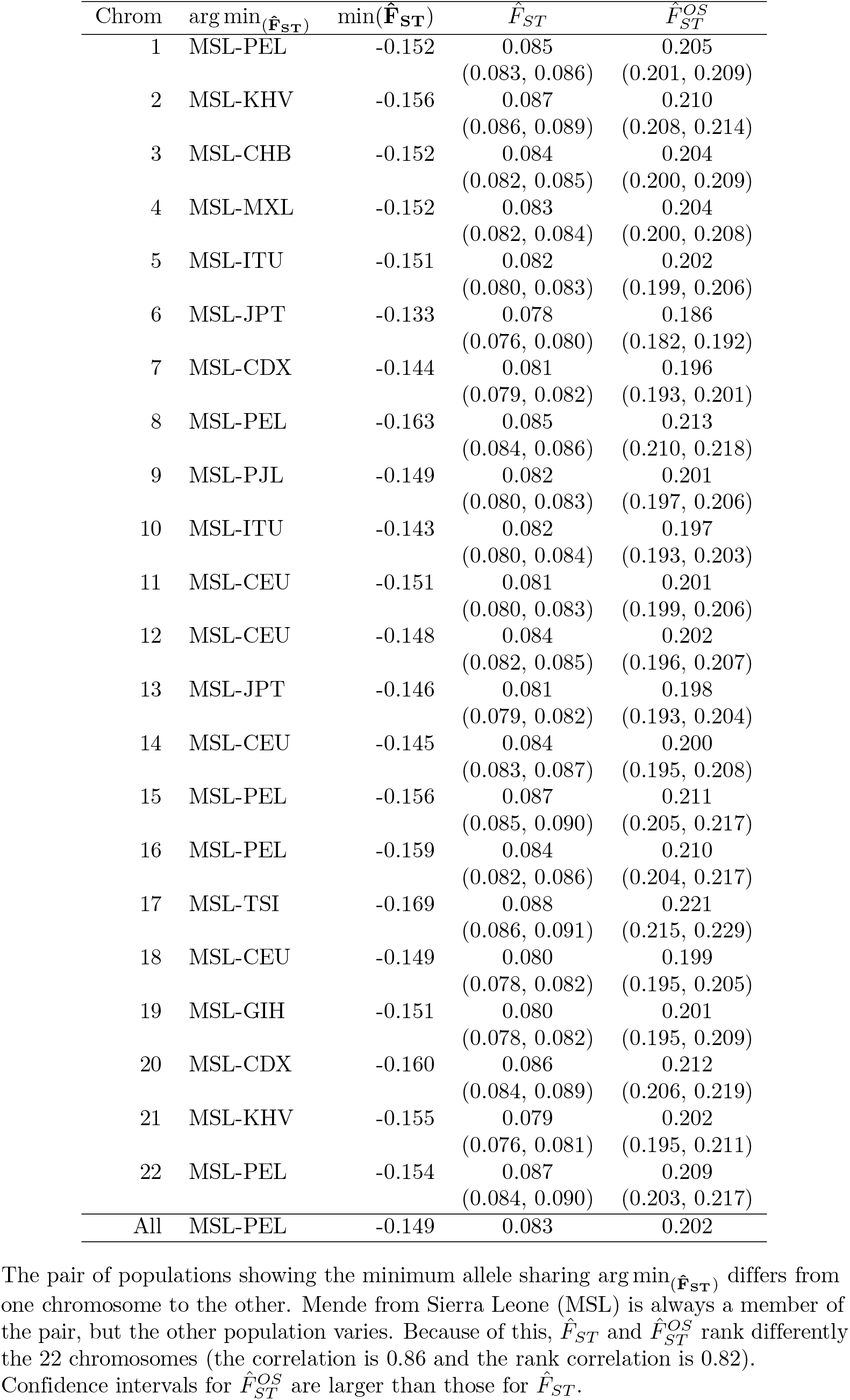
Comparison of 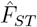 (the average of population specific 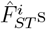, Eq. 8) and 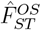 in the 1000 genomes data set for each chromosome. Confidence intervals obtained by block bootstrap, using 100kb blocks The pair of populations showing the minimum allele sharing arg 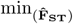 differs from one chromosome to the other. Mende from Sierra Leone (MSL) is always a member of the pair, but the other population varies. Because of this, 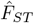 and 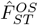 ST rank differently the 22 chromosomes (the correlation is 0:86 and the rank correlation is 0:82). Confidence intervals for 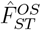 ST are larger than those for 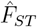.

**Fig S3.**
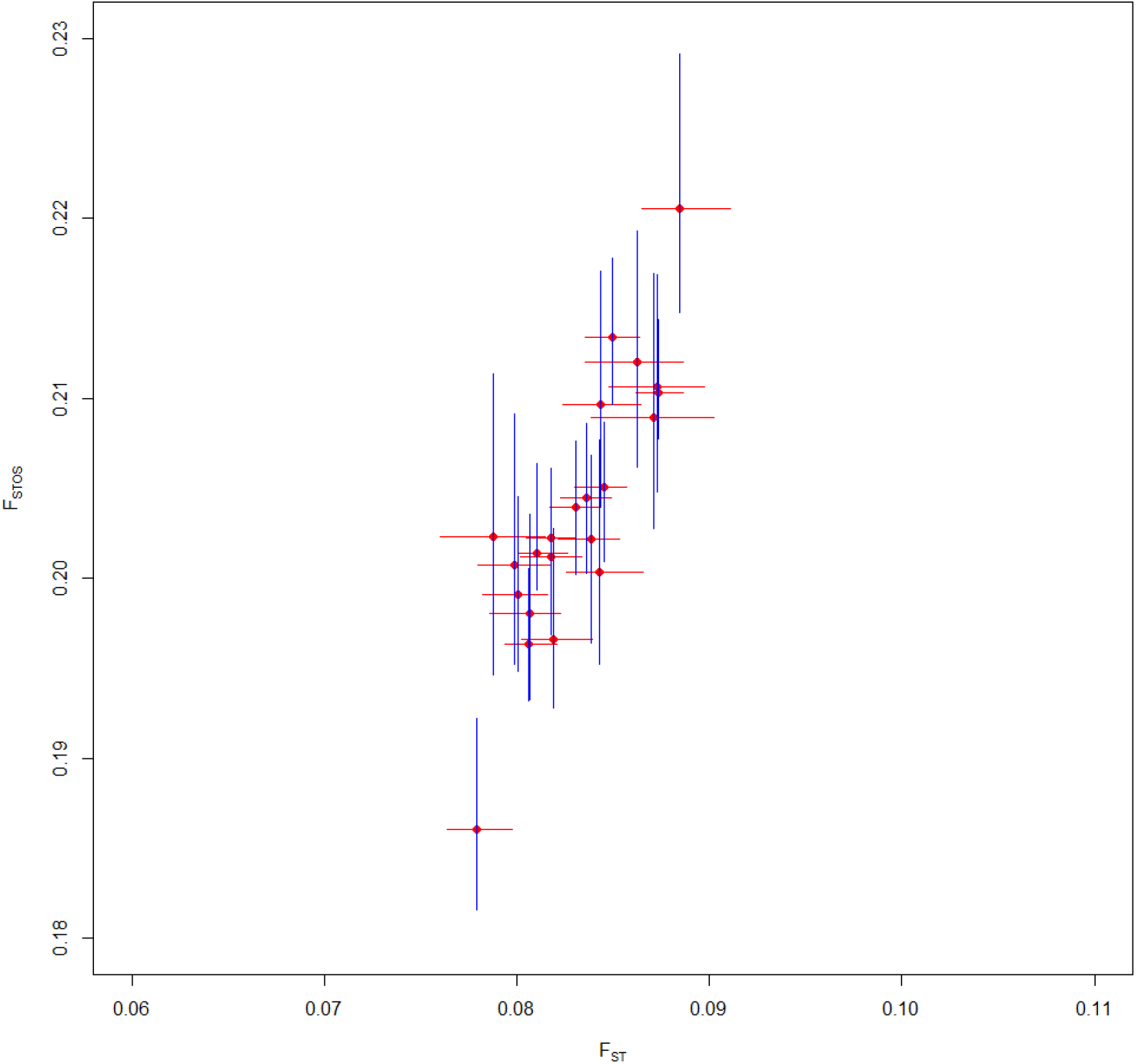
chromosome specific 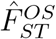 against 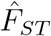 from the 1000 genomes data and their block bootstrap confidence intervals (CI). Note the x- and y-axis cover the same *F*_*ST*_ interval width (0.05). CIs for 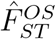 in blue are between 2.25 and 3.6 times wider than CIs for 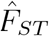 in red.

